# Increased functional connectivity of the intraparietal sulcus underlies the attenuation of numerosity estimations for self-generated words

**DOI:** 10.1101/2020.12.18.423390

**Authors:** Giedre Stripeikyte, Michael Pereira, Giulio Rognini, Jevita Potheegadoo, Olaf Blanke, Nathan Faivre

**Affiliations:** Center for Neuroprosthetics, Swiss Federal Institute of Technology (EPFL), CH-1202 Geneva, Switzerland; Brain Mind Institute, Faculty of Life Sciences, Swiss Federal Institute of Technology (EPFL), CH-1015 Lausanne, Switzerland; Univ. Grenoble Alpes, Univ. Savoie Mont Blanc, CNRS, LPNC, 38000 Grenoble, France; Department of Neurology, University of Geneva, CH-1211 Geneva, Switzerland

## Abstract

Previous studies have shown that self-generated stimuli in auditory, visual, and somatosensory domains are attenuated, producing decreased behavioral and neural responses compared to the same stimuli that are externally generated. Yet, whether such attenuation also occurs for higher-level cognitive functions beyond sensorimotor processing remains unknown. In this study, we assessed whether cognitive functions such as numerosity estimations are subject to attenuation. We designed a task allowing the controlled comparison of numerosity estimations for self (active condition) and externally (passive condition) generated words. Our behavioral results showed a larger underestimation of self-compared to externally-generated words, suggesting that numerosity estimations for self-generated words are attenuated. Moreover, the linear relationship between the reported and actual number of words was stronger for self-generated words, although the ability to track errors about numerosity estimations was similar across conditions. Neuroimaging results revealed that numerosity underestimation involved increased functional connectivity between the right intraparietal sulcus and an extended network (bilateral supplementary motor area, left inferior parietal lobule and left superior temporal gyrus) when estimating the number of self vs. externally generated words. We interpret our results in light of two models of attenuation and discuss their perceptual versus cognitive origins.

## INTRODUCTION

The ability to distinguish self-versus externally-generated stimuli is crucial for self-representation (Kircher and David, 2003; Legrand, 2006). A typical mechanism by which stimuli generated by oneself and those caused by external sources are distinguished is sensory attenuation, whereby self-generated stimuli are perceived as less intense. Indeed, previous studies have shown that self-produced stimuli in the auditory (Baess et al., 2011; Timm et al., 2014), visual (Hughes and Waszak, 2011; Benazet et al., 2016) and somatosensory domains (Shergill et al., 2013; Kilteni and Ehrsson, 2020), are perceived as less intense compared to the same stimuli when they are externally generated. Such attenuation was shown at the behavioral level and at the neural level in sensory cortical regions (e.g., auditory cortex (Rummell et al., 2016; Whitford, 2019)) as well as the thalamus, cerebellum, supplementary motor area and inferior parietal cortex (Hickok, 2012; Lima et al., 2016; Bansal et al., 2018; Brooks and Cullen, 2019).

Previous studies have shown that attenuation not only applies to stimuli that are generated by overt actions, but also extends to covert processes such as motor imagery (Kilteni et al., 2018) or inner speech (Scott et al., 2013; Whitford et al., 2017). In the latter case, attenuation of the electrophysiological (Whitford et al., 2017; Jack et al., 2019) and behavioral (Scott et al., 2013) responses corresponding to the test stimulus (phoneme) was observed when it matched the imagined cued phoneme. These studies used phonemes that are integrated into the early stages of hierarchical speech processing implying primary sensory cortices (Liebenthal et al., 2005; DeWitt and Rauschecker, 2012), where attenuation effects have been demonstrated (Rummell et al., 2016). An outstanding question is whether attenuation of self-generated stimuli is limited to sensory consequences (e.g., weakened percept) or whether it can also impact cognitive processes that are non-perceptual in nature such as abstract judgments or performance monitoring.

To answer this question, we investigated the cognitive function of numerosity estimations, defined as approximate judgments when counting is not involved (Dehaene, 1997). Properties of numerosity estimations such as innateness, amodality, or precision that linearly decreases with increasing numerosity have been described (Anobile et al., 2016; Burr et al., 2018). Extensive neuroimaging work has also established that brain areas in the intraparietal sulcus (IPS) region play a key role in numerosity processing (for review, see Arsalidou & Taylor, 2011). If attenuation for self-related functions extends to higher-level cognitive processes such as numerosity estimations, we would expect the number of items that are self-generated to be underestimated compared to externally-generated items in relation to modulation of IPS activity. Thus, this study aimed at investigating whether numerosity estimations of self-generated words were attenuated compared to numerosity estimations of passively heard words, thereby demonstrating that self-attenuation applies to cognitive processes beyond sensory processing.

To investigate which brain regions and networks showed activity related to attenuation processes, we designed a functional magnetic resonance imaging (fMRI) paradigm allowing the controlled comparison of numerosity estimations of either self-generated words (active condition) during a phonetic verbal fluency task or externally-generated words (passive condition) while passively listening to a stream of words. Additionally, we asked participants to evaluate the error they could have made in their estimation (performance monitoring). Assuming that attenuation occurs during numerosity estimations (active condition), we predicted that participants would underestimate the number of self-vs. externally-generated words and explored possible effects on performance monitoring. As performance monitoring is better for self-initiated vs. observed processes (Pereira et al., 2020), we expected to find a stronger relationship between the reported and actual number of words in the active vs. passive condition. At the neural level, considering that sensory attenuation of self-generated stimuli involves the corresponding sensory brain regions (e.g., primary auditory cortex for attenuated sounds, see Rummell et al., 2016; Whitford, 2019), we expected to find reduced BOLD signal during the active vs. passive condition in areas responsible for numerosity processing including the IPS.

## METHODS

### Participants

Two independent participant groups were tested in this study. First, we performed a behavioral pilot experiment in a mock scanner, where we tested 17 (6 women) participants (age range: 18 – 28 years; M = 23 years, CI(95%) = [23, 24]); schooling level varied between 13 and 21 years; M = 17 years, CI(95%) = [16, 18]). During the main fMRI experiment, we studied 25 (14 women) participants (age range: 18 – 37 years; M = 23 years, CI(95%) = [22, 26]; schooling level varied from 12 to 22 years: M = 17 years, CI(95%) = [16, 18]). All participants were right-handed according to the Edinburgh Hand Preference Inventory (Oldfield, 1971) and native French-speaking healthy volunteers with no history of neurological or psychiatric disease and no recent reported history of drug use. Participants had normal or corrected-to-normal vision and no claustrophobia. All participants were naive to the purpose of the study, gave informed consent in accordance with institutional guidelines and the Declaration of Helsinki, and received monetary compensation (20 CHF / hour). The study was approved by the local ethical committee of the canton Geneva (protocol ID: 2015-00092).

### Experimental task

The experiment was performed in a mock scanner environment (both for the behavioral pilot experiment and the training for the main fMRI experiment) and in the MRI scanner (main fMRI experiment). The task consisted in phonetic verbal fluency and passive word listening parts, followed by numerosity and error estimations (Figure 1). The task comprised two conditions (20 trials each) during which participants either covertly generated words (active condition) or listened to pre-recorded words (passive condition). Each trial started with a randomly jittered inter-trial interval varying between 4 s and 4.75 s, followed by an audio cue (2 s) indicating which of the active or passive condition will follow, and a recorded cue letter (2 s). Next, the word generation phase lasted between 20 s and 35 s; in the active condition, participants had to covertly generate words starting with the cued letter (Figure 1, top). For each word they covertly generated, participants were asked to press a response button, which gave us access to the actual number of generated words. In the passive condition, a series of pre-recorded audio words were played to the participants, who were asked to press the response button for every word that contained the cue letter (Figure 1, bottom). In both conditions, the end of the generation/listening phase was indicated by an audio cue (0.5 s). After that, participants first reported the estimation of a total number of words generated (active condition) or heard (passive condition). For this, they used two buttons that moved a slider displayed on the screen. The slider was presented as a random integer (ranging from 0 to 20), which could be changed in value by pressing the response box buttons (left button – decrease the value, right button – increase the value). This was followed by an error estimation, where participants were asked to evaluate their performance on the numerosity estimation by estimating the magnitude of the error they thought they might have made (in number of words). For this, an automatically sliding bar was presented, and the participant had to select with one button press the desired value (e.g., +/-2 words error). Values varied from +/-0 words (e.g., numerosity response judged as correct) to +/-5 words mistaken. The total time for numerosity and error estimations were restricted to a maximum of 7 s. Except for the numerosity and error estimation periods, participants were asked to perform the task with eyes closed.

**Figure 1.**
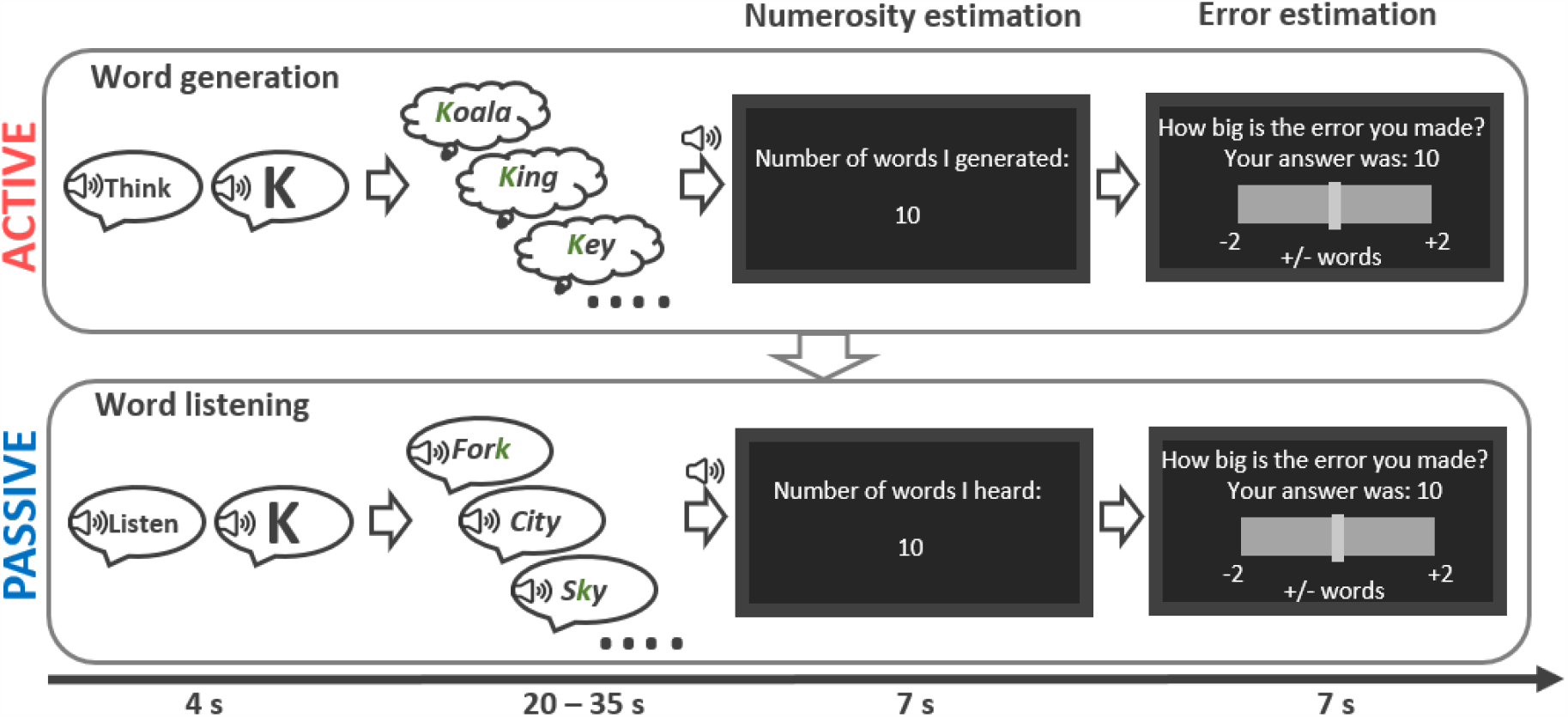
Schematic representation of the general task flow: active (top) and passive (bottom) conditions consisting of instruction, word generation/listening, numerosity and error estimation. Each condition was repeated 4 times per block. Auditory and visual instructions were provided in French.

### Stimuli

All stimuli were prepared and presented using Matlab 2016b (mathworks.com) and the Psychtoolbox-3 toolbox (psychtoolbox.org; Brainard 1997; Kleiner et al. 2007; Pelli 1997). Twenty different cue letters were used for active and passive conditions. The same cue letter was used once during the active and once during the passive condition in the counterbalanced order. Played back words during the passive condition were chosen from a list of 420 French words (Ferrand and Alario, 1998). The audio stimuli presented during the task were recorded by male and female native French speakers in a neutral manner and registered in *wav* format with 11025Hz sampling frequency. The gender of the voice pronouncing words in the passive condition was matched to the participants’ gender. During the experiment, participants were equipped with MRI compatible earphones and report buttons for the right hand.

### Procedure

Participants were trained in a mock scanner prior to the main fMRI experiment in order to familiarize themselves with the task. They were asked to perform four trials of the task during the training (twice for each condition), with the cue letters ‘j’ and ‘k’. These letters were not used later during the main fMRI experiment.

The main experiment consisted of three runs lasting approximately 15 min, each with short breaks in-between. The total duration of a trial varied between 40 and 55 s due to pseudorandomized time for the word generation/ listening phase. This time variability was introduced to avoid habituation and predictability for the number of generated/heard words and to decorrelate hemodynamic activity related to word generation/listening and numerosity estimations. The experiment was designed in 10 blocks with 4 trials of the same condition per block. The blocks of the active condition trials always preceded the blocks of the passive condition, allowing us to use the number and pace of generated words that were recorded based on participant’s button presses to playback the words during the next block of the passive condition. The order of the number of words played during the passive condition was shuffled within the block. This was done to ensure that participants could not recognize whether the number and pace of the played words matched the preceding active condition block.

After the main experiment, a standard phonemic verbal fluency (generation time of 60 s, cue letter ‘p’) test (Lezak, 1995) was performed overtly to verify that subjects understood the task correctly. Overall, the experiment lasted approximately 1h 30 min (MRI session) and 1h in the mock scanner (pilot session). The pilot mock scanner study contained the same procedure as the main MRI experiment, except for the shorter breaks between the runs since there was no scanning involved.

### Behavioral performance measures

Most statistical frameworks to analyze performance monitoring have been developed for discrimination tasks with a binary response (e.g., Fleming and Lau, 2014 for a review). In the following, we propose two indices of numerosity performance and performance monitoring to analyze ordinal data (e.g., number of words). In addition to the prerequisites described below, these indices were defined at the single-trial level so they could serve as parametric regressors of interest in the fMRI analysis.

The numerosity performance index is an accuracy ratio reflecting how correctly participants estimated the number of words during the generation/listening phase. For each trial, we wanted signed numerosity performance to be proportional to the difference between the reported number of generated/heard words (numerosity estimation; [N]) and the actual number of words generated/heard [W]. We normalized this difference by the sum of numerosity estimation and actually generated/heard number of words [N+W] (equation 1) to give more weight to errors made about low numbers of words (e.g., an error of +/-2 given a numerosity estimation of 8 has higher magnitude than an error of +/-2 given a numerosity estimation of 16; Figure 1-2 A). Negative numerosity performance values thus reflected an underestimation of generated/heard words, and positive values reflected an overestimation of generated/heard words. In contrast, null numerosity performance values reflected correct answers about the number of generated/heard words.

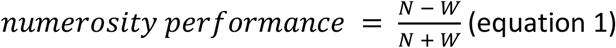

Performance monitoring reflected how well participants estimated an error about their previous performance. We defined it as the absolute value of the difference between the error estimation [E] and (accuracy [N-W]), normalized by the sum of the numerosity estimation and words generated/heard [N+W]) (equation 2). Normalization was done in order to consider the difficulty; the same error made when estimating the low number of words or high number of words should be penalized proportionally. A performance monitoring value of 0 reflected ideal error tracking, whereby participants correctly estimated the error made during the numerosity estimation. An increase in performance monitoring value represented an increase in error magnitude while estimating the difference between numerosity estimation and the actual number of words generated/heard (Figure 1-2 B, C).

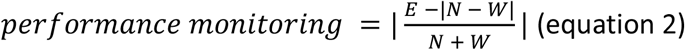

Trials for which participants did not generate at least five words or failed to answer numerosity or error estimations within the time limit were excluded from behavioral and fMRI analysis (in total, 2.44 ± 1.87 trials/subject were excluded). The threshold of 5 words was selected according to the working memory capacity of 5 ± 2 items (Cowan, 2010). Considering all participants’ data, 6.1% of all trials were discarded.

### Behavioral data analysis

All continuous variables (numerosity performance, performance monitoring) were analyzed using linear mixed-effects regressions with condition (“active”, “passive”) as a fixed effect and a random intercept by participant and condition. The inclusion of additional random effects was guided by model comparison and selection based on Bayesian Information Criteria. Analyzes were performed using the lme4 (Bates et al., 2015) and lmerTest (Kuznetsova et al., 2017) packages in R (R-project.org). The significance of fixed effects was estimated using Satterthwaite’s approximation for degrees of freedom of F statistics (Luke, 2017).

### fMRI data acquisition

MRI data were acquired using a Siemens Magnetom Prisma 3 T scanner with a 64-channel head coil. T1 weighted (1 mm isotropic) scans were acquired using a Magnetization Prepared Rapid Acquisition Gradient Echo (MPRAGE) sequence (TR = 2300 ms; TI = 900 ms; TE = 2.25 ms; flip-angle = 8 degrees; GRAPPA = 2; FOV = 256 × 256 mm; 208 slices).

Functional scans were obtained using echo-planar (EPI) sequence (multiband acceleration = 6; TR = 1000 ms; TE = 32 ms; flip-angle = 58 degrees; FOV = 224 × 224 mm; matrix = 64 × 64; slice thickness = 2 mm; number of slices = 66). The number of functional image volumes varied according to the experiment duration (2278 ± 61 volumes).

### fMRI data preprocessing

Anatomical and functional images were processed and analyzed using SPM-12 (Welcome Department of Cognitive Neurology, London, UK). Pre-processing steps included slice time correction, field-map distortion correction, realignment and unwarping to spatially correct for head motions and distortions, co-registration of structural and functional images, normalization of all images to common Montreal Neurological Institute (MNI) space, and spatial smoothing with a Gaussian kernel with a full-width at half maximum (FWHM) of 4 mm. Quality assurance of all EPI images was performed with the criteria of maximum 2 mm translation and 2° rotation between volumes. In addition, an excessive movement was estimated with the mean framewise displacement (FD) (Power et al., 2012) with the exclusion threshold of 0.5 mm. None of the subjects had a higher mean FD (0.2 ± 0.06 mm) than the set threshold.

### fMRI data analysis

We used a two-level random-effects analysis. In the first-level analysis, condition-specific effects were estimated according to a general linear model (GLM) fitted for each subject. An average mask of grey matter from all subjects was built using FSL (fsl.fmrib.ox.ac.uk/fsl) and used to mask out white matter and non-brain tissues. The GLM was built using six boxcar regressors corresponding to the duration of the word generation/listening phase, numerosity estimation and error estimation in the active and passive conditions. Parametric modulators of numerosity performance and performance monitoring were included in the numerosity estimation and error estimation regressors, respectively. Further, we added regressors of no interest corresponding to audio instructions, button presses and excluded trials, plus six regressors for head motion (translation and rotation).

At the second-level (group level), we performed a one-way analysis of variance (ANOVA) with F-tests to assess main effects common to active and passive conditions and t-tests to analyze the difference between conditions (active vs. passive) for each regressor of interest: numerosity estimation and error estimation. We used a voxel-level statistical threshold of p<0.001 and corrected for multiple comparisons at the cluster level using family-wise error (FWE) correction with the threshold of p<0.05. We used the anatomical automatic labelling (AAL) atlas for brain parcellation (Tzourio-Mazoyer et al., 2002).

### Functional connectivity analysis

Psychophysiological Interaction (PPI) analysis was employed to identify modulations of functional coupling between a seed region and other brain regions by experimental conditions (active vs. passive) (Friston et al., 1997). To perform this analysis we used generalized psychophysiological interaction (gPPI) toolbox version 13.1 (McLaren et al., 2012). Spheres of 6 mm radius were formed around the peak coordinates of the right IPS (x = 29; y = -65; z = 50) and the left IPS (x = -27; y = -66; z = 47) clusters that were identified in the second-level analyses. First-level (individual) GLM analyses were performed, including task regressors of numerosity estimation or error estimation (psychological term) and time course of the seed region (physiological term). As for other aforementioned fMRI analyses, we performed t-tests to compare the differences between conditions.

### Data and code availability

The MRI data that support the findings of this study will be uploaded on a public repository upon publication. Behavioral data, analysis and task codes are available here: https://gitlab.epfl.ch/lnco-public/cognitive-attenuation.

## RESULTS

### Behavioral results

By design, the number of generated and heard words was matched between conditions (active: M = 10.7, SD = 3.6, CI(95%) = [10.4, 11.1]; passive: M = 10.7, SD = 3.5, CI(95%) =[10.4, 11.0], F_(1,884)_ = 0.005, p = 0.94). The duration of word generation and listening did not differ between conditions either (active: M = 27.4, SD = 5.6, CI(95%) = [26.9, 27.9]; passive: M = 28.4, SD = 7.0, CI(95%) = [27.7, 29.0], F_(1,26)_ = 2.04, p = 0.16). Having shown that conditions were similar in terms of difficulty, we turned to the analysis of numerosity performance and performance monitoring.

Numerosity performance indices revealed that globally, participants underestimated the number of words (M = -0.05, SD = 0.11, CI(95%) = [-0.06, -0.05]). This underestimation was significantly larger (F_(1,24)_ = 5.85, p = 0.023, 18 subjects out of 25 showed the effect) in the active condition (M = -0.07, SD = 0.11, CI(95%) = [-0.08, -0.06]) compared to the passive condition (M = -0.04, SD = 0.12, CI(95%) = [-0.05, -0.03]) (Figure 2 A). Comparable results were obtained during the pilot experiment in the mock scanner in an independent group of subjects (active: M = -0.05, SD = 0.01, CI(95%) = [-0.06, -0.03]); passive: M = -0.03, SD = 0.10, CI(95%) = [-0.04, -0.02]; F_(1,16)_ = 5.80 p = 0.016). These two experiments suggest that numerosity estimations for self-generated words are attenuated.

**Figure 2.**
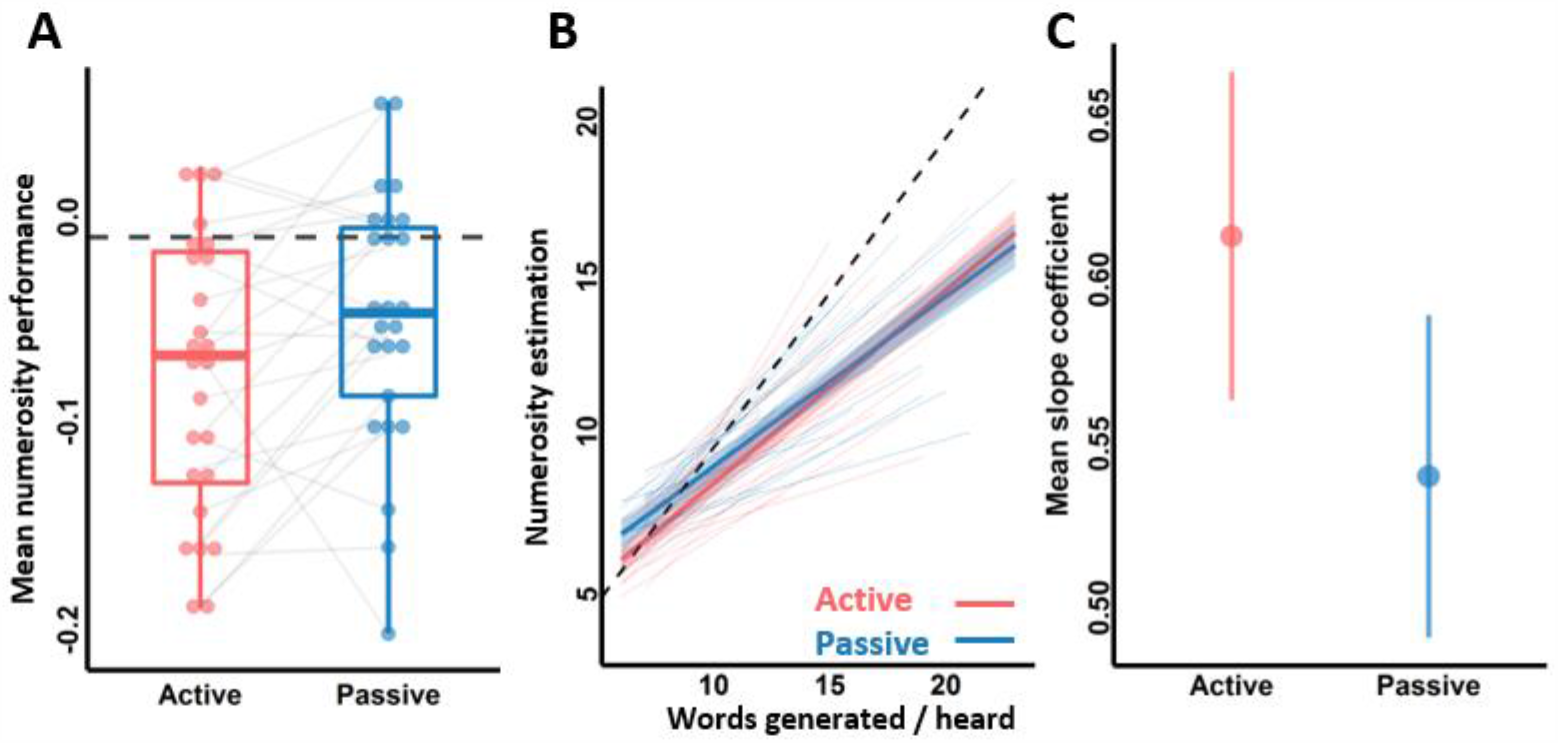
Behavioral results. (A) Individual numerosity performance in the active (red) and passive (blue) conditions. A value of zero represents ideal performance, where the reported and actual number of words match. Each dot represents a participant mean numerosity performance. Grey lines connect performance from the same participant across conditions (B) Mixed-effects linear regression between numerosity estimation (reported number of words) and an actual number of words. Reference dashed line with a slope equal to 1 represents ideal performance. Thick lines represent the global model fit, while thin lines depict individual estimates. (C) Mean slope coefficient estimates with 2.5% and 97.5% confidence intervals.

We then assessed the linear relationship between the reported (numerosity estimation) and actual number of words (Figure 2 B). Besides a main effect of words (F_(1,908)_ = 1042, p < 0.001) and of condition (F_(1,198)_ = 8.25, p = 0.0045), we found an interaction between condition and generated/heard words (F_(1,802)_ = 4.3, p = 0.038). This interaction was driven by a steeper slope in the active condition (M = 0.62, CI(95%) = [0.57, 0.67]) compared to the passive condition (M = 0.55, CI(95%) = [0.50, 0.60]) (Figure 2 C), which indicates better numerosity tracking for self-generated words during the main fMRI experiment (a slope of value 1 reflecting ideal performance). We note that this interaction did not reach significance in our pilot experiment (F_(1,350)_ = 1.10, p = 0.66), which calls for interpreting this result with caution.

Next, we investigated whether performance monitoring varied between conditions, by quantifying how well participants were able to track the error made when estimating the number of generated/heard words. We found no differences in performance monitoring (F_(1,24)_ = 1.09, p = 0.3) between the active (M = 0.07, SD = 0.06, CI(95%) = [0.07, 0.08]) and passive (M = 0.08, SD = 0.06, CI(95%) = [0.07, 0.09]) conditions during the main fMRI experiment, nor during the mock scanner pilot experiment (F_(1,16)_ = 0.78, p = 0.39; active: M = 0.07, SD =0.06, CI(95%) = [0.06, 0.08]; passive: M = 0.06, SD =0.06, CI(95%) = [0.06, 0.07]). This confirms the absence of evidence supporting an effect of attenuation on performance monitoring.

Finally, we carried out a control task for word generation outside the scanner, during which participants had to perform a standard verbal fluency task overtly with the cue letter ‘p’. Comparing the number of words generated, starting with the letter ‘p’ overtly (outside the scanner) and covertly (during the scanning), we did not observe any significant differences between the number of words generated overtly (M = 12.6, SD = 3.79, CI(95%) = [10.9, 14.1]) and covertly (M = 12.3, SD = 3.96, CI(95%) = [10.6, 13.9]; paired t-test; t_(24)_ = -0.4; p=0.69). This control task confirms that subjects did generate words covertly and comparably to overt fluency, as instructed.

### fMRI results

#### Numerosity performance

Brain activity during the numerosity estimation phase was widespread, irrespective of the experimental condition (Table 1-1). Differences between conditions revealed widespread relative deactivations in the active compared to the passive condition in the bilateral parietal cortex including IPS, middle-superior temporal gyri, precuneus, cerebellum, middle cingulate gyri, SMA, insula, middle-superior frontal gyri, hippocampus, caudate nucleus, putamen (for the detailed list of all areas see Table 1-2).

To avoid contaminating the results with inherent differences between the active and passive conditions that are not specific to numerosity processes, we looked for brain activity parametrically modulated by numerosity performance. We found a single brain region with such a pattern of activity, namely the right IPS, whose activity was negatively correlated with numerosity performance (main effect F = 20.0, p_FWE_ = 0.042; Figure 3; Table 1). The same pattern was found bilaterally when using a less stringent threshold (p<0.005 peak level uncorrected; Table 1-3, Figure 3-1). Unlike our prediction, IPS activation did not differ between the active and passive conditions (p_FWE_ > 0.05), suggesting that this effect was independent of whether words were heard or actively generated.

**Table 1.**
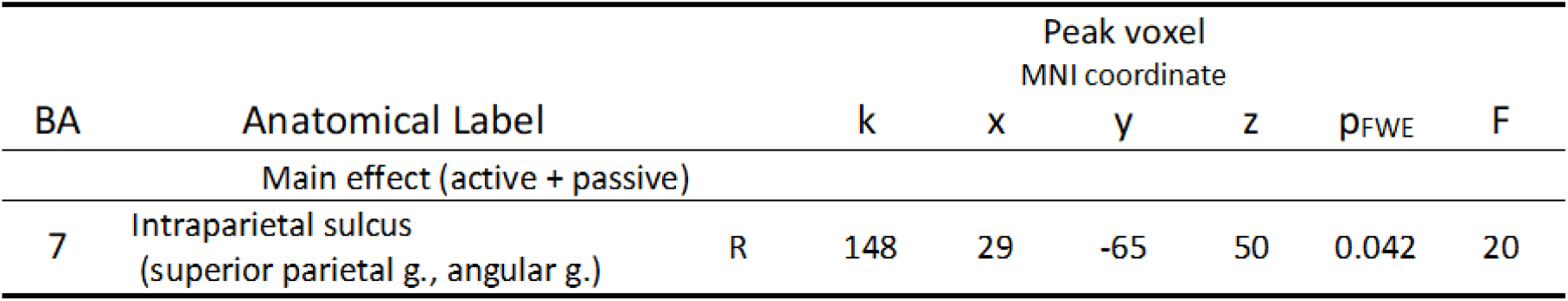
Parametric modulation of numerosity performance. Brain areas with BOLD signal correlating with trial-by-trial behavioral measures of numerosity performance during numerosity estimation. Voxel level p<0.001 uncorrected, cluster threshold at p<0.05 FWE corrected. BA – Broadmann area, k – cluster size, R – right hemisphere

**Figure 3.**
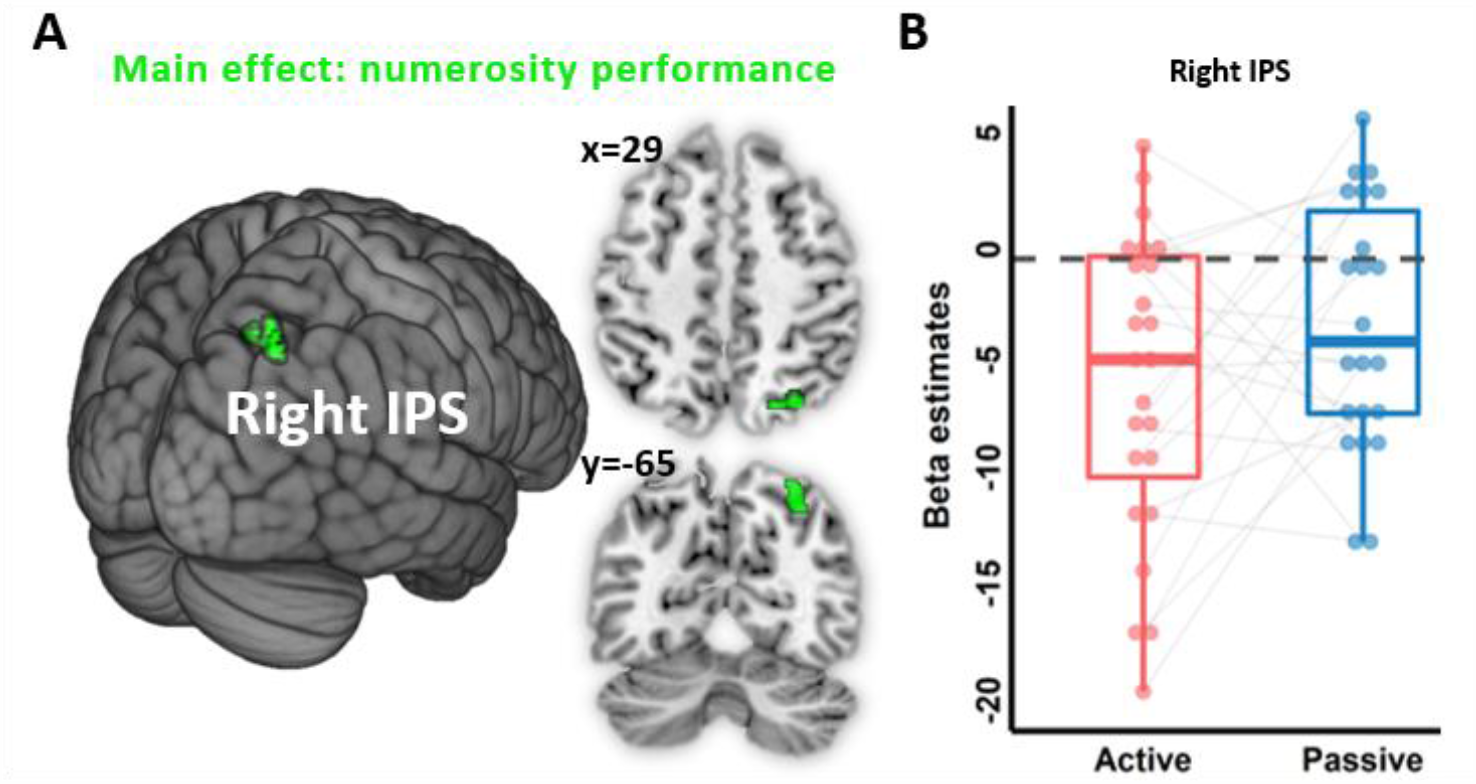
Neural correlates of numerosity performance. (A) Main effect (active+passive conditions) of the parametric modulation of numerosity performance during numerosity estimation in the right intraparietal sulcus (IPS, depicted in green). (B) Box plot of the corresponding individual beta estimates in the right IPS. Each dot represents a participant mean beta estimates. Grey lines connect beta estimates from the same participant across conditions.

Using PPI analysis, we investigated whether the functional connectivity of the right IPS with other brain regions differed between the active and passive conditions. This analysis revealed that bilateral supplementary motor area (SMA), left inferior parietal lobule (IPL) and left superior temporal gyrus (STG) had increased connectivity with the right IPS in the active compared to the passive condition (Figure 4, Table 2). No effect in the other direction was observed.

**Figure 4.**
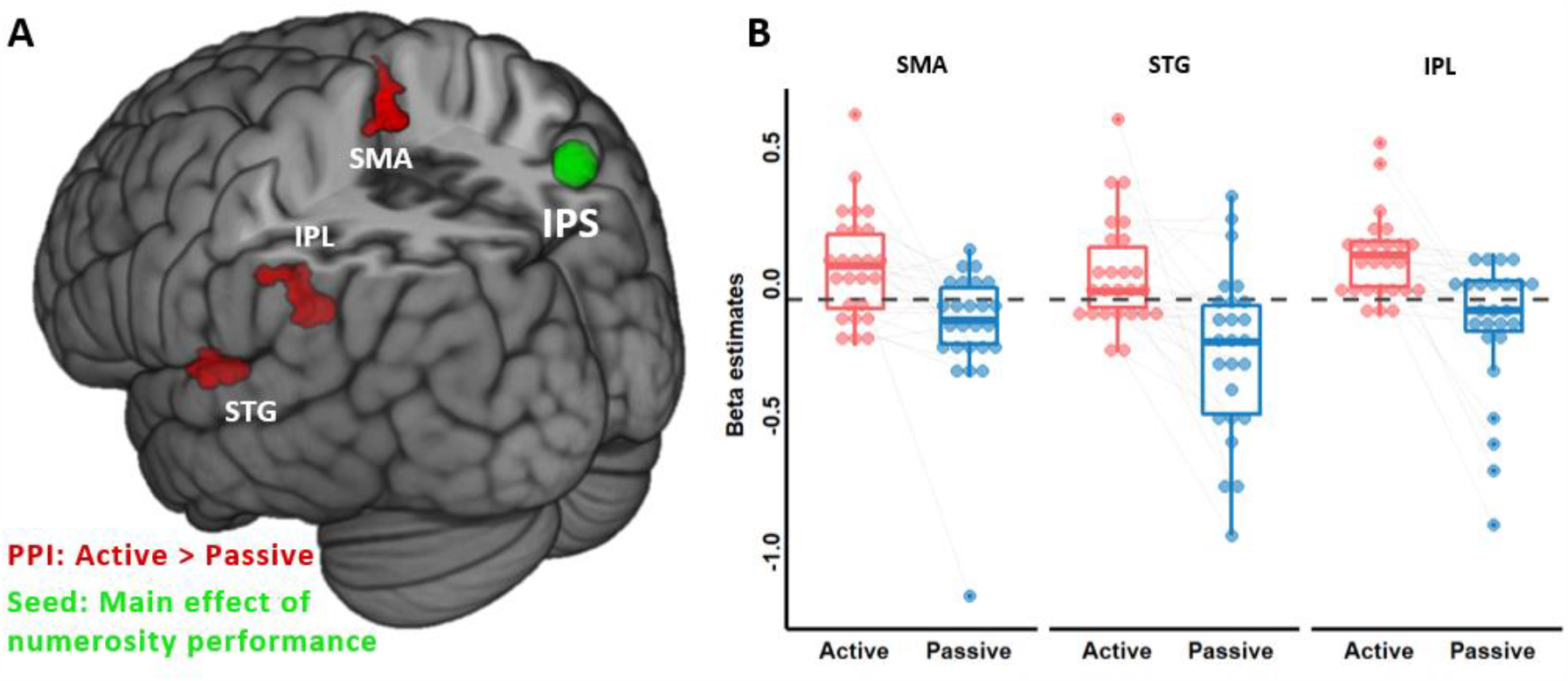
PPI analysis: numerosity estimation. (A) Increased functional connectivity in the active vs. passive condition between the right intraparietal sulcus seed (IPS, depicted in green) and bilateral supplementary motor area (SMA), left inferior parietal lobule (IPL) and superior temporal gyri (STG), depicted in red. (C) Box plots of the corresponding beta estimates for each significant PPI cluster. Each dot represents a participant mean beta estimates. Grey lines connect beta estimates from the same participant across conditions.

**Table 2.** PPI analysis: numerosity estimation. Brain areas identified through PPI analysis seeding from the right intraparietal sulcus during numerosity estimation. Voxel level p<0.001 uncorrected, cluster threshold at p<0.05 FWE corrected. BA – Broadmann area, k – cluster size, R – right hemisphere, L – left hemisphere

| BA | Anatomical Label |  | k | Peak voxel<br>MNI coordinate |  |  | p<sub>FWE</sub> | T |
| --- | --- | --- | --- | --- | --- | --- | --- | --- |
|  |  |  |  | x | y | z |  |  |
| Active > passive |  |  |  |  |  |  |  |  |
| 41<br>42 | Superior temporal g. | L | 177 | -54 | -20 | 6 | 0.038 | 5.14 |
| 40 | Supramarginal g.,<br>parietal inferior g. | L | 291 | -44 | -42 | 29 | 0.003 | 5.14 |
| 6 | Paracentral lobule, SMA | L/R | 167 | -3 | -30 | 63 | 0.049 | 4.59 |

### Performance monitoring

Besides numerosity performance, we also quantified brain activity during the error estimation phase. We found widespread cortical and subcortical activations during this phase compared to baseline, irrespective of the experimental condition (Table 3-1).

Moreover, bilateral insula and right putamen were significantly less active during the active compared to the passive condition (Figure 5; Table 3). The left caudate nucleus showed the opposite pattern, with higher activity in the passive than in the active condition.

**Figure 5.**
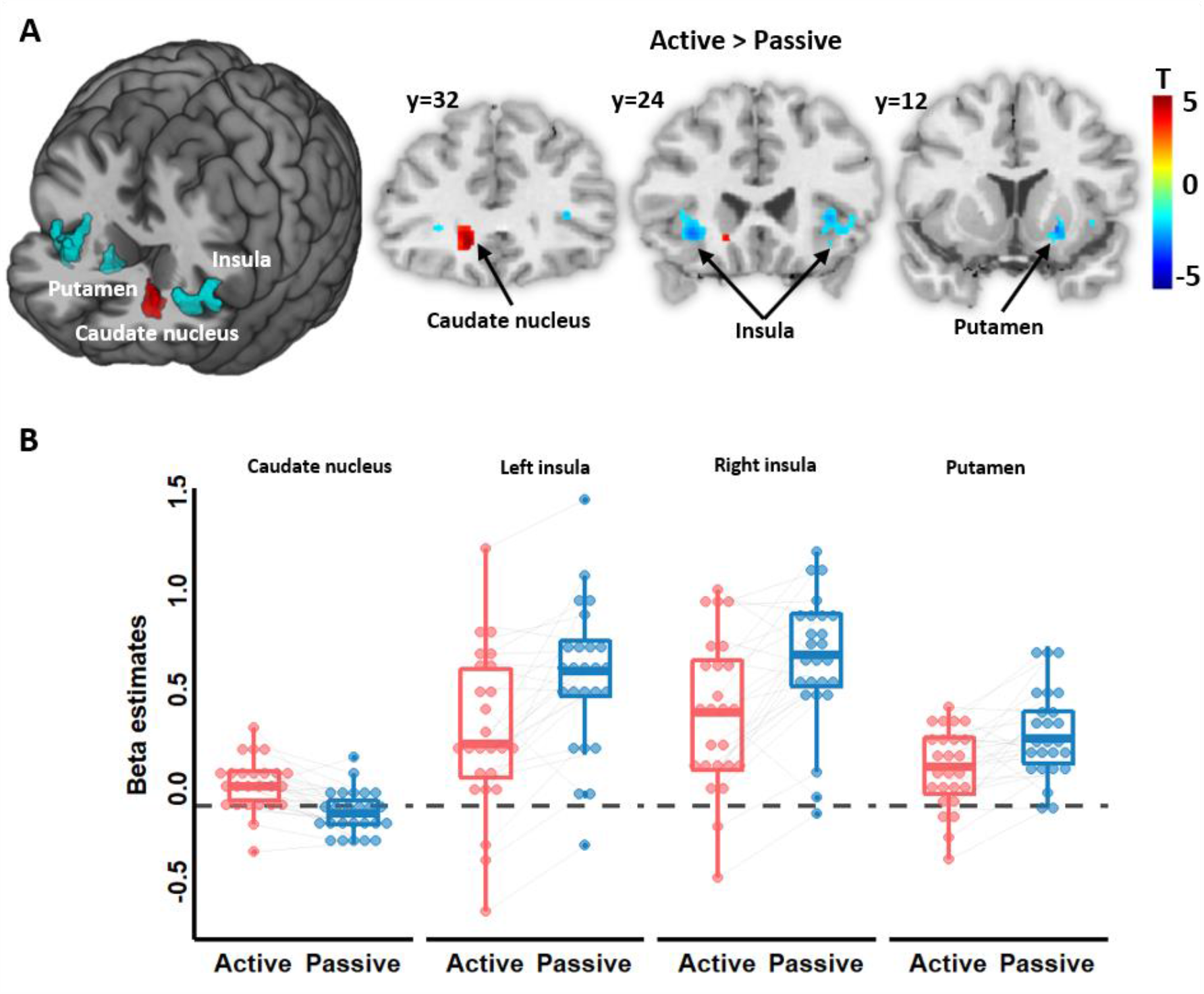
Neural correlates of error estimations. (A) Decreased neural activity during the active vs. passive condition in the right putamen and insula bilaterally (depicted in blue). Increased activity in the active vs. passive condition in the left caudate nucleus (depicted in red). (B) Box plots of the corresponding beta estimates in the active (red) and passive (blue) conditions for each significant cluster observed. Each dot represents a participant mean beta estimates. Grey lines connect beta estimates from the same participant across conditions.

**Table 3.**
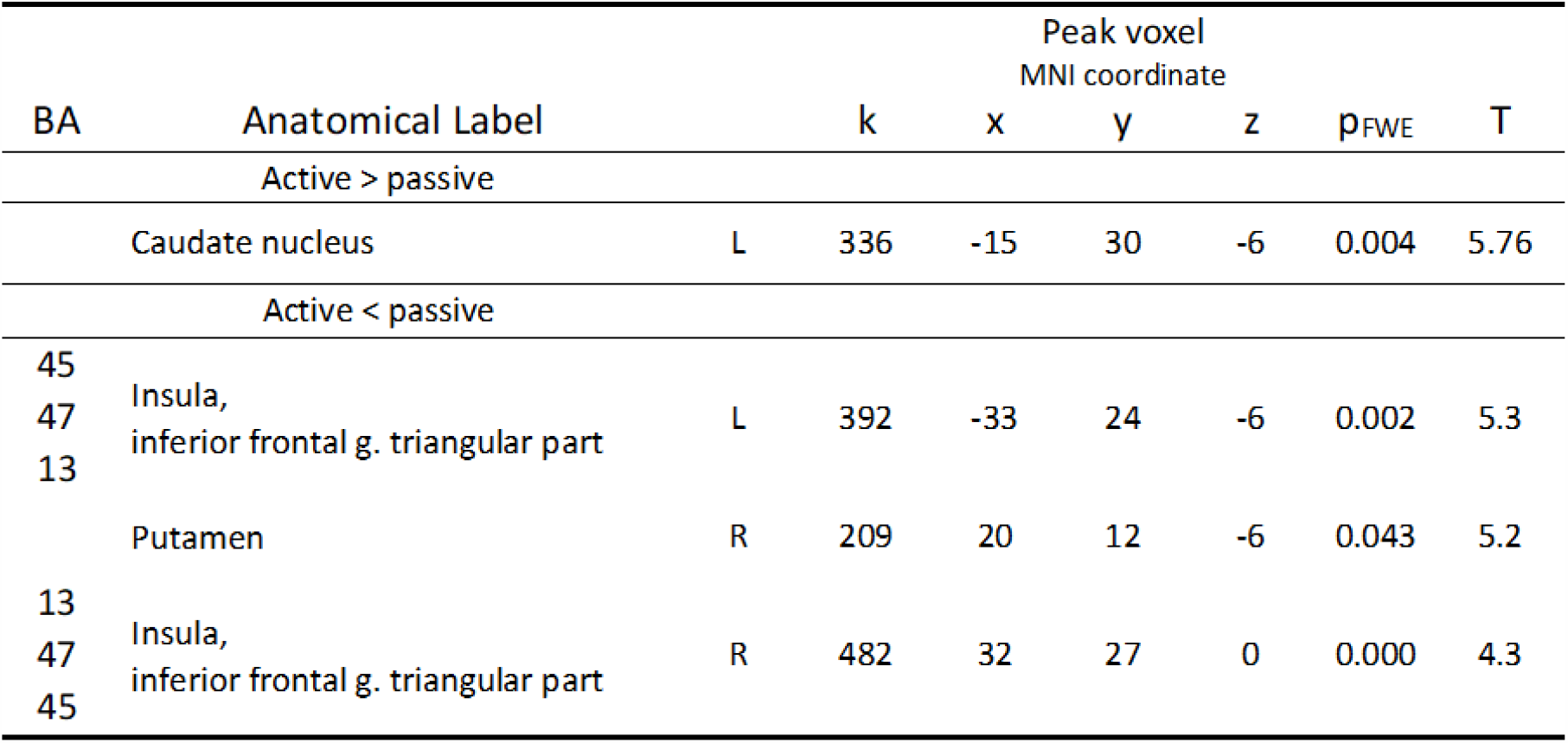
Brain activations during error estimations. Voxel level p<0.001 uncorrected, cluster threshold at p<0.05 FWE corrected. BA – Broadmann area, k – cluster size, R – right hemisphere, L – left hemisphere

To avoid confounding factors between the active and passive condition, we looked more specifically at parametric modulations of error monitoring. We observed that BOLD signal in the left IPS was more related to performance monitoring in the active compared to the passive condition (t = 4.02, p_FWE_ = 0.041; Figure 6; Table 4). Interestingly, this activation cluster overlapped with the main effect of numerosity performance (see Figure 3-1).

**Figure 6.**
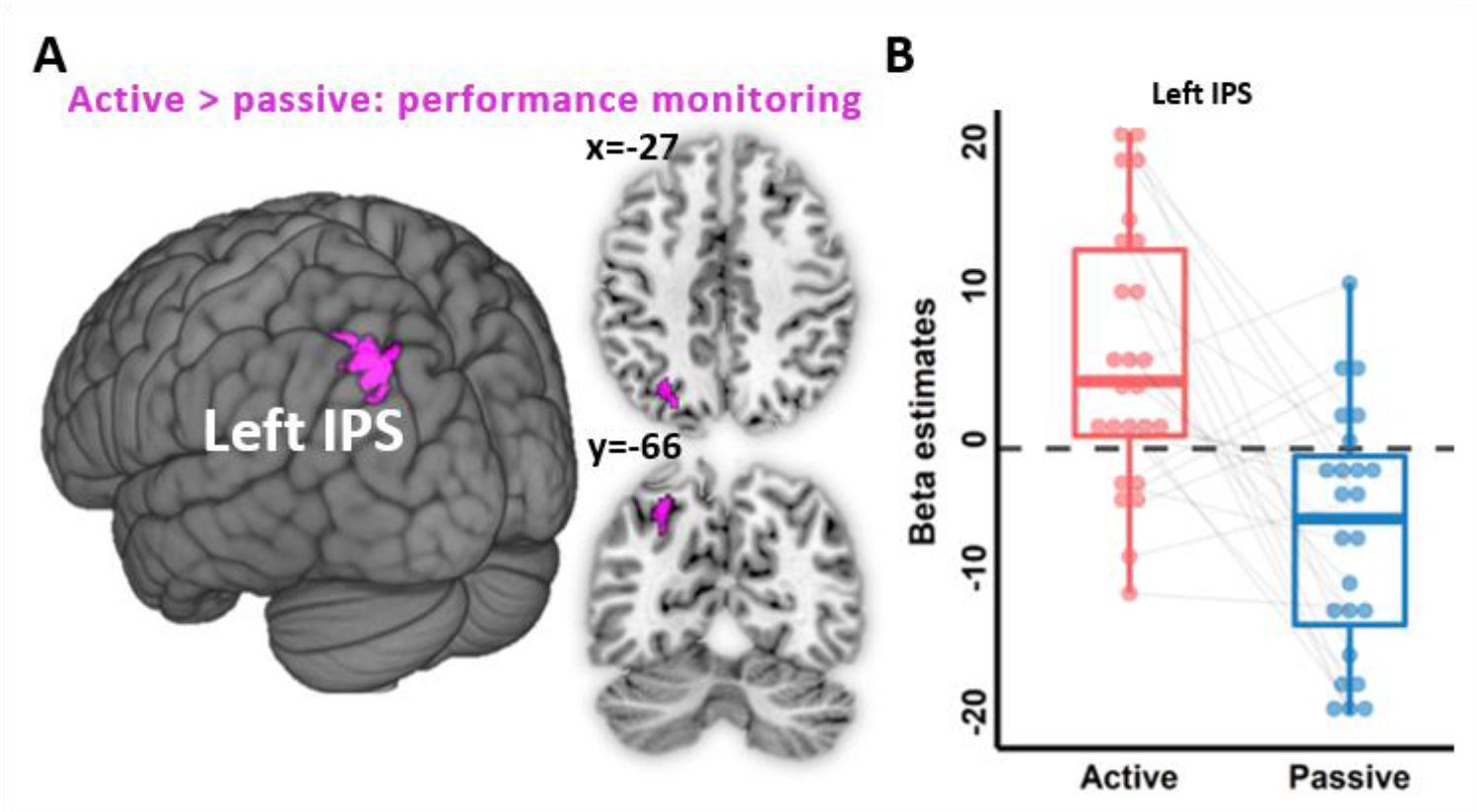
Parametric modulation of performance monitoring. (A) Positive association between the trial-wise performance monitoring and BOLD signal during performance monitoring in the active vs. passive condition in the left intraparietal sulcus (IPS, depicted in purple). (B) Box plot of corresponding beta estimates in the left IPS in the active (blue) and passive (red) conditions. Each dot represents a participant mean beta estimates. Grey lines connect beta estimates from the same participant across conditions.

**Table 4.** Parametric modulation of performance monitoring. Brain areas with BOLD signal correlating with trial-by-trial behavioral measures of performance monitoring during error estimation. Voxel level p<0.001 uncorrected, cluster threshold at p<0.05 FWE corrected. BA – Broadmann area, k – cluster size, L – left hemisphere

| BA | Anatomical Label |  | k | Peak voxel<br>MNI coordinate |  |  | p<sub>FWE</sub> | T |
| --- | --- | --- | --- | --- | --- | --- | --- | --- |
|  |  |  |  | x | y | z |  |  |
| Active > passive |  |  |  |  |  |  |  |  |
| 7 | Intraparietal sulcus<br>(superior/inferior parietal g.) | L | 198 | -27 | -66 | 47 | 0.041 | 4.02 |

Lastly, results from the PPI analysis using the left IPS as a seed ROI did not reveal any functional connectivity differences between the active and passive conditions during error estimation.

## DISCUSSION

The present study examined whether attenuation of self-generated stimuli impacts cognitive processes beyond the traditionally investigated sensory processes, namely numerosity estimations and performance monitoring. To this end, we developed an experimental paradigm allowing the controlled comparison of numerosity estimations and performance monitoring regarding self- and externally-generated words while acquiring fMRI data. We found that participants more strongly underestimated the number of self-compared to externally-generated words, providing behavioral evidence that numerosity estimations are indeed subject to attenuation. As expected, numerosity performance was associated with hemodynamic activity changes in IPS. Furthermore, a network including the bilateral SMA, left IPL and left STG showed increased functional connectivity with the right IPS during numerosity estimations for self-generated words, suggesting that numerosity-related attenuation involves this neural network. Finally, by asking participants to monitor the accuracy of their own numerosity estimations, we found equivalent performance monitoring for self- and externally-generated words. Although no difference was found at the behavioral level, we found that performance monitoring was associated with increased IPS activity in the active vs. passive condition.

### Behavioral and neural markers of attenuated numerosity estimations

Participants underestimated the number of words they generated or heard. Such underestimations were previously described regarding the numeric estimation of perceptual quantities (e.g., number of dots or sequences of sounds) to discrete measures (e.g., Arabic numeral) (Castronovo and Seron, 2007; Reinert et al., 2019). In the present study, we found that word numerosity underestimation was stronger in the active compared to the passive condition, in line with our hypothesis that attenuation of self-generated stimuli may extend to higher-level cognitive functions (Kilteni et al., 2018; Jack et al., 2019).

Attenuation of self-generated stimuli has been mostly investigated for sensory processes and refers to the diminished behavioral and neural responses associated with self-generated compared to externally-generated stimuli (Timm et al., 2014; Benazet et al., 2016). Importantly, attenuation has recently also been described for imagined actions in the absence of overt actions (e.g., Kilteni et al., 2018; Jack et al., 2019;). For example, imagined self-touch was felt as less intense compared to externally applied touch (Kilteni et al., 2018). Similarly, imagined speech elicited reduced electrophysiological signals related to auditory processing (Whitford et al., 2017). These studies, however, investigated cognitive processes intimately linked to sensorimotor systems (e.g., imagined movement and touch), yet, to the best of our knowledge, it was unknown whether comparable mechanisms of attenuation also affect cognition beyond sensorimotor processing, such as numerosity estimations with no or less obviously implicated sensorimotor processes (e.g., Dehaene, 1997). Here we show that differential attenuation can be observed for cognitive processes such as numerosity estimations that depend on whether they are self-generated or not. One leading theory of sensory attenuation of self-generated stimuli is the internal forward model (Farrer and Frith, 2002; Miall and Wolpert, 1996), based on the proposal that corollary discharges related to action are used to predict the sensory consequences of that action. When such predictions match the actual sensory feedback from the action, its sensory consequences are attenuated, and the action is perceived as self-generated (Wolpert and Flanagan, 2001). The forward model thus proposes to link sensory attenuation to the sensory predictions generated by a neural comparator. An analogous mechanism has been proposed to account for attenuation for covert actions such as motor imagery (Kilteni et al., 2018) or inner speech (Tian and Poeppel, 2010). The present data show that attenuation exists beyond overt and covert actions, raising the possibility that the forward model extends to repetitive cognitive activity (e.g., fluency and related numerosity estimations). One possibility, similar to what has been described for inner speech, would be that numerosity underestimation stems from a weaker perceptual representation of self-generated words. In other words, attenuation may not impact numerosity estimations directly, but rather through a decreased representation of self-generated words, which in turn may lead to attenuation of numerosity estimations. A second account could be that mental operations have a gating effect (Cromwell et al., 2008), independent from predictive mechanisms, thereby directly affecting the strength of the mental representations of self-generated words. A similar gating mechanism has recently been shown to offer a plausible alternative to forward model accounts of sensory self-attenuation (Thomas et al., 2020).

At the neural level, the IPS showed activity related to numerosity performance, in line with previous studies (Cohen and Dehaene, 1996; Piazza et al., 2006). This relation, however, was not modulated by condition (active vs. passive), as one could have expected based on previous reports showing that sensory attenuation coincides with decreased activity in the corresponding sensory area (e.g., attenuation in primary auditory cortex during self-generated auditory stimuli; Rummell et al., 2016; Whitford, 2019). Importantly, we found differences in functional coupling during numerosity estimations of the IPS with a network of brain regions between the active and passive conditions. The increase in coupling during the active condition occurred between the IPS and a network comprising the SMA, IPL and STG, known to be involved in predictive processing, for example of self-generated auditory and imagery speech (Lima et al., 2016; Tian et al., 2016). Although previous neuroimaging studies have mainly shown attenuated brain activity for self-generated actions (e.g., Whitford, 2019), increases in functional connectivity such as we describe have been reported in the primary auditory cortex during attenuated speech (van de Ven et al., 2009), or of somatosensory cortex during attenuated touch (Kilteni and Ehrsson, 2020). Thus, increased functional connectivity in a network centered on the key numerosity region, IPS, and that has been associated with speech-related processing further supports that numerosity underestimation stems from a process related to attenuation when participants self-generate words.

Besides numerosity underestimation, we observed that participants formed more accurate numerosity judgments for self-vs. externally-generated words: while participants’ word estimations were lower in the active condition than in the passive condition, the relation between their estimated number of words and the actual number of words was better in the active condition vs. passive condition, suggesting a sharper representation of the number of self-generated words. This result corroborates recent findings showing better monitoring for decisions that are committed rather than observed (Pereira et al., 2020). This improved monitoring of self-generated words could be related to the sharpening of expected representations known in the sensory domain (Kok et al., 2012) or to a self-generation effect underlying the facilitation of information encoding and enhanced recall for self-generated stimuli (Bertsch et al., 2007; Slamecka and Graf, 1978).

### Performance monitoring is modulated at the neural but not behavioral level

Previous research has shown that both humans and non-human primates (Beran et al., 2006; Duyan and Balcı, 2019, 2018) can monitor the quality of their numerosity estimations. Thus, in addition to asking participants to estimate the number of words they generated or heard, we also asked them to estimate their own error during this process (performance monitoring). Although our behavioral results showed similar performance monitoring between conditions, we observed that performance monitoring was associated with hemodynamic activity in the left IPS predominantly in the active condition. Interestingly, this region is not typically associated with performance monitoring such as the prefrontal cortex or the insula/inferior frontal gyrus (Vaccaro and Fleming, 2018). Since activity in the left IPS is related to numerosity estimation (Cappelletti et al., 2007; Dormal et al., 2012), this parametric modulation could therefore represent a substrate for monitoring specific to numerosity estimations. By investigating global fMRI activity during error estimations, we also observed decreased activation in the insula and putamen and increased activity in the caudate nucleus when comparing the active vs. passive conditions. To note, the anterior insula and putamen were activated during both conditions but were attenuated in the active compared to the passive conditions. While the anterior insula activations are consistent with previous literature on performance monitoring (Ullsperger et al., 2010; Bastin et al., 2017), the findings of modulated activity in the striatum were unexpected.

Both areas are known as essential for the control of goal-directed decision-making (Balleine et al., 2007; Kim and Im, 2019), suggesting that different activations of these regions between conditions could have acted as an alternative cognitive mechanism impacting similar performance monitoring.

## Conclusion

Based on behavioral and neuroimaging data, we propose that higher-level cognitive functions such as numerosity estimations about the number of self-generated words are attenuated. Such attenuation involves a functional network including a key-numerosity region (IPS) and speech-related regions including the SMA, IPL and STG. While attenuating the sensory consequences of one’s actions is of crucial importance for aspects of the self, such as the sense of agency, attenuating the products of one’s mental activities may also be relevant to distinguish them from external sources of information. Our paradigm offers a promising tool to investigate attenuation processes related to the self in cognition and to compare and distinguish them from sensory attenuation processes. It may be of relevance to the study of clinical cases in which attenuation in sensory and cognitive domains may be is altered, including patients with psychotic symptoms like thought insertion whereby thoughts are not considered as one’s own, but of somebody else.

## Acknowledgments

The authors are grateful for the technical support during data acquisition to the Foundation Campus Biotech Geneva (FCBG) Human Neuroscience Platform of the Campus Biotech. This work was supported by the Bertarelli Foundation (grant number 532024), the Swiss National Science Foundation (grant number 3100A0-112493), National Center of Competence in Research (NCCR) “Synapsy - The Synaptic Bases of Mental Diseases” (grant number 51NF40 – 185897), and two generous donors advised by Carigest SA. Nathan Faivre has received funding from the European Research Council (ERC) under the European Union Horizon 2020 research and innovation programme (grant number 803122).

## Author Contributions

GS, NF, GR, OB developed the study concept and contributed to the study design. Testing, data collection and data analysis were performed by GS. GS, MP, JP and NF provided methodological support. GS, NF, MP, and OB drafted the paper; all authors provided critical revisions and approved the final version of the paper for submission.

