## Extended data for "Increased functional connectivity of the intraparietal sulcus underlies the attenuation of numerosity estimations for self-generated words"

**Abbreviated title:** Attenuation of numerosity estimations


**Authors**
Giedre Stripeikyte[1,2], Michael Pereira[1,2,3], Giulio Rognini[1,2], Jevita Potheegadoo[1,2], Olaf Blanke[1,2,4]*, Nathan Faivre[1,2,3]*

**Affiliations**
1. Center for Neuroprosthetics, Swiss Federal Institute of Technology (EPFL), CH-1202 Geneva, Switzerland
2. Brain Mind Institute, Faculty of Life Sciences, Swiss Federal Institute of Technology (EPFL), CH-1015 Lausanne, Switzerland
3. Univ. Grenoble Alpes, Univ. Savoie Mont Blanc, CNRS, LPNC, 38000 Grenoble, France
4. Department of Neurology, University of Geneva, CH-1211 Geneva, Switzerland

* These authors contributed equally to this study

**Corresponding authors**
Olaf Blanke
Bertarelli Chair in Cognitive Neuroprosthetics
Center for Neuroprosthetics & Brain Mind Institute
School of Life Sciences
Campus Biotech
Swiss Federal Institute of Technology
Ecole Polytechnique Fédérale de Lausanne (EPFL)
CH – 1202 Geneva


Nathan Faivre
Laboratoire de Psychologie et Neurocognition
CNRS UMR 5105, UGA BSHM
1251 Avenue Centrale, 38058 Grenoble Cedex 9, France


**Data simulation of derived measures of numerosity performance and performance monitoring**

We simulated data to assess the validity of our indices of numerosity performance and performance monitoring. Data was simulated with pseudorandom generation of an actual number of words and numerosity estimations ranging between 5 and 20. Error estimation varied between 0 and 5 as set during the task design. Simulated data showed that our derived formula accounting for the underestimation and overestimation of numerosity performance changes linearly (Figure 1-1 A). The formula accounting for the performance monitoring considers the difficulty of the task. In other words, for the bigger number words, the performance monitoring value will not be as high as for a smaller number of words even though the subject chose the same value for the error estimation (Figure 1-1 B).

**FIGURES AND TABLES**

**Extended figures**

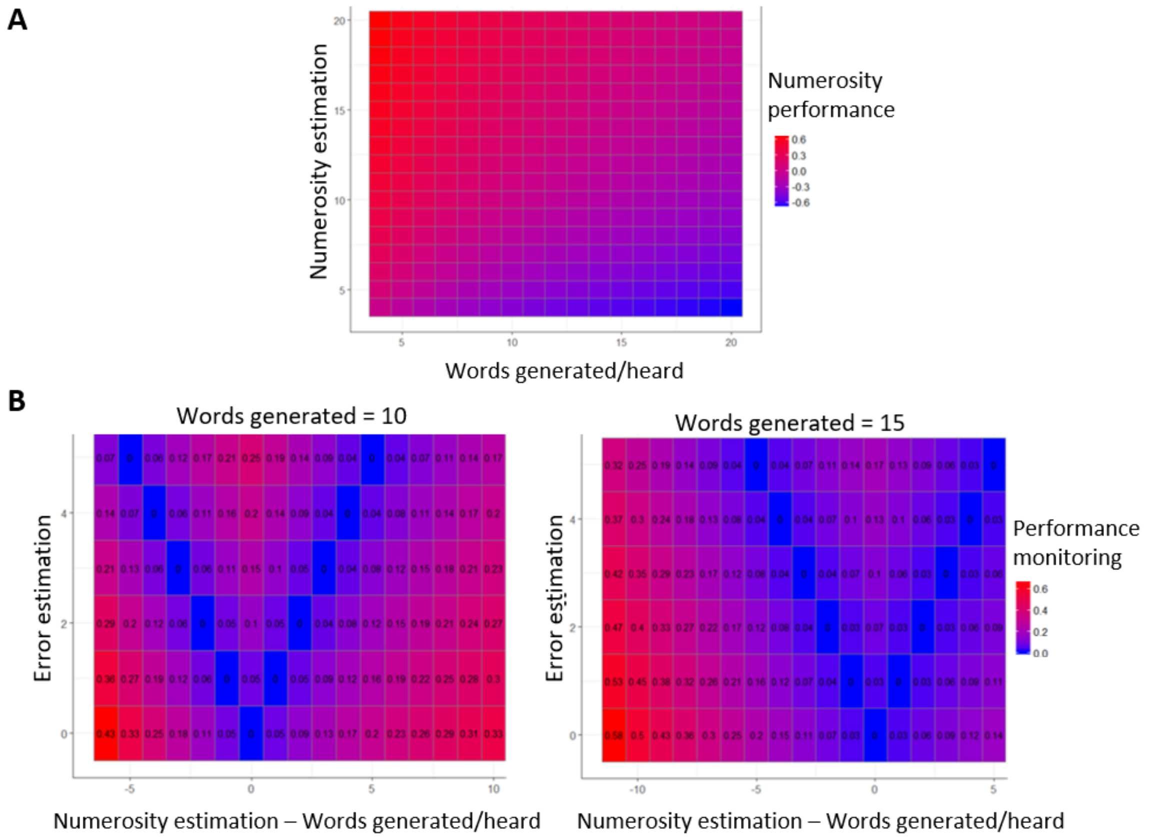


**Figure 1-1.** Data simulation of behavioral performance measures. (A) Simulated numerosity performance. Increasing intensity of red represent linear increase of overestimation (positive values of numerosity performance), while increasing intensity in blue represent linear increase of underestimation (negative values of numerosity performance). (B) Simulated performance monitoring when the number of words is constant (10 or 15 words). The performance monitoring value is depicted in the heatmap changing from blue (being correct) to red (worse performance monitoring).

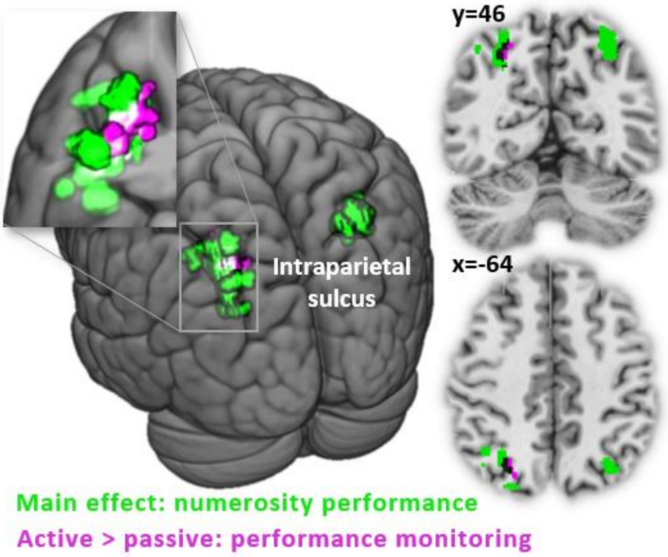


**Figure 3-1.** Overlap between the main effect (active + passive conditions) of numerosity performance during numerosity estimation (depicted in green) and parametric modulation of performance monitoring during the error report (active vs. passive condition) (depicted in purple); peak level uncorrected $p<0.001$. peak level uncorrected $p<0.005$.

### Extended tables

| BA | Anatomical Label | | k | Peak voxel MNI coordinate x | y | z | $p_{FWE}$ | F |
|---|---|---|---|---|---|---|---|---|
| | Main effect (active + passive) | | | | | | | |
| 7<br>18<br>40<br>19<br>17<br>31<br>30<br>37<br>2<br>23<br>20<br>21<br>29<br>3 | Parietal inferior, superior g.<br>Occipital inferior, middle, superior g.<br>Cerebellum<br>Lingual g.<br>Calcarine<br>Cuneus<br>Precuneus<br>Fusiform g.<br>Temporal inferior, middle g.<br>Angular g.<br>Postcentral g.<br>Supramarginal g. | L/R | 88149 | 6 | -74 | 14 | <0.001 | 261.5 |
| 6<br>9<br>8<br>10<br>32<br>13<br>46<br>44 | Frontal inferior, middle, superior g.<br>Precentral g.<br>SMA<br>Caudate<br>Cingulate anterior, middle, posterior g.<br>Insula<br>Precentral g.<br>Thalamus<br>Hippocampus<br>Putamen | L/R | 56151 | 24 | -27 | -2 | <0.001 | 232.4 |
| 13<br>21<br>22<br>40<br>6<br>41 | Rolandic opercular<br>Temporal middle, superior g.<br>Insula<br>Hippocamus | R | 4107 | 44 | -9 | 0 | <0.001 | 74.9 |
| | Corpus callosum | | 740 | 17 | -33 | 20 | <0.001 | 58.7 |
| 10<br>11<br>32 | Orbital part of medial frontal g.<br>Cingulum anterior | L/R | 2565 | -9 | 41 | -14 | <0.001 | 50.3 |
| 21<br>13 | Temporal middle, superior g.<br>Hippocampus<br>Insula<br>Amygdala | L | 2558 | -24 | -12 | -14 | <0.001 | 47.8 |
| 39 | Angular g. | L | 563 | -30 | -74 | 26 | <0.001 | 43.3 |
| 4 | Precentral g. | R | 832 | 36 | -21 | 54 | <0.001 | 42.2 |
| | Hippocampus<br>Corpus callosum | L | 1074 | -20 | -47 | 15 | <0.001 | 41.9 |
| 13 | Temporal superior g. | L | 176 | -47 | -29 | 20 | 0.044 | 21.3 |

**Table 1-1. Numerosity estimation: main effect (active + passive conditions).** Brain areas activated during the numerosity estimation phase independent of parametric modulation. Voxel level p<0.001 uncorrected, cluster threshold at p<0.05 FWE corrected. BA –

Broadmann area, k – cluster size, R – right hemisphere, L – left hemisphere, R – right hemisphere

| | | | | Peak voxel MNI coordinate | | | | |
|---|---|---|---|---|---|---|---|---|
| BA | Anatomical Label | | k | x | y | z | $p_{FWE}$ | T |
| | Active < passive | | | | | | | |
| 7<br>6<br>40<br>31<br>13<br>24<br>21<br>4<br>32<br>39<br>3<br>5<br>9<br>23<br>27<br>8<br>18<br>44 | Middle temporal g.<br>Precuneus<br>Superior temporal g.<br>Postcentral<br>Middle cingulum<br>Cerebellum<br>SMA<br>Insula<br>Precentral<br>Supramarginal g.<br>Angular g.<br>Superior parietal<br>Putamen<br>Middle frontal g.<br>Caudate<br>Superior fontal g.<br>Superior parietal g.<br>Calcarine<br>Rolandic percular<br>Supramarginal g.<br>Thalamus<br>Hippocampus<br>Fusiform g.<br>Inferior temporal g.<br>Lingual g.<br>Cuneus<br>Occipital g. | L/R | 113587 | 20 | 8 | -12 | | 9.75 |
| | Thalamus | L | 314 | -11 | -23 | 15 | | 6.84 |
| 10 | Middle frontal g. | L | 442 | -36 | 50 | 27 | | 4.97 |
| 10 | Middle frontal g. | R | 399 | 29 | 53 | 32 | | 4.69 |

**Table 1-2. Numerosity estimation: difference between active and passive conditions.** Brain areas showing a difference in activity between the active and passive condition during the numerosity estimation phase, independent of parametric modulation. Voxel level p<0.001 uncorrected, cluster threshold at p<0.05 FWE corrected. BA – Broadmann area, k – cluster size, R – right hemisphere, L – left hemisphere, R – right hemisphere

| BA | Anatomical Label | | k | Peak voxel MNI coordinate x | y | z | $p_{FWE}$ | T |
|---|---|---|---|---|---|---|---|---|
| | Main effect (active + passive) | | | | | | | |
| 7 | Intraparietal sulcus (superior parietal g., angular g.) | R | 466 | 29 | -65 | 50 | 0.004 | 20 |
| 7 | Intraparietal sulcus (superior/inferior parietal g., middle occipital g.) | L | 758 | -30 | -63 | 42 | 0.000 | 15.9 |

**Table 1-3. Parametric modulation of numerosity performance.** Brain areas in which the BOLD signal correlates with the trial-by-trial behavioral measures of numerosity performance during the numerosity estimation phase. Voxel level p<0.005 uncorrected, cluster threshold at p<0.05 FWE corrected. BA – Broadmann area, k – cluster size, R – right hemisphere, L – left hemisphere

| BA | Anatomical Label | | k | Peak voxel MNI coordinate | | | $p_{FWE}$ | F |
|---|---|---|---|---|---|---|---|---|
| | | | | x | y | z | | |
| | Main effect (active + passive) | | | | | | | |
| 7<br>18<br>19<br>6<br>40<br>9<br>8<br>37<br>10<br>32<br>31<br>17<br>30<br>23<br>2<br>46<br>13<br>39 | Frontal inferior, middle, superior g.<br>Supramarginal g.<br>Parietal inferior, superior g.<br>Occipital inferior, middle, superior g.<br>Cerebellum<br>Lingual g.<br>Calcarine<br>Cuneus<br>Precuneus<br>Caudate nucleus<br>Putamen<br>Thalamus<br>Fusiform g.<br>Temporal inferior, middle g.<br>Angular g.<br>Postcentral g.<br>Supramarginal g.<br>SMA<br>Cingulate anterior, middle g.<br>Insula<br>Hippocampus | L/R | 174684 | 2 | -75 | 8 | <0.001 | 462.4 |
| | Corpus callosum<br>Precuneus | R | 1177 | 23 | -44 | 14 | <0.001 | 134.1 |
| | Hippocampus | L | 224 | -26 | -12 | 14 | 0.014 | 58.9 |
| 38 | Pole of temporal superior g.<br>Insula | R | 362 | 44 | 0 | -11 | 0.001 | 56.4 |
| 38 | Pole of temporal superior g.<br>Insula | L | 513 | -44 | 2 | -15 | <0.001 | 55.4 |
| 11<br>32<br>24 | Orbital part of medial frontal g.<br>Cingulum anterior g. | L/R | 835 | 3 | 24 | -5 | <0.001 | 44.9 |
| 13 | Insula | R | 264 | 39 | -15 | 9 | 0.006 | 34.6 |
| 21 | Temporal middle g. | L | 196 | -56 | -12 | -15 | 0.027 | 28.6 |

**Table 3-1. Error estimation main effect (active + passive conditions).** Brain areas with activations during the error estimation phase, independent of parametric modulation. Voxel level p<0.001 uncorrected, cluster threshold at p<0.05 FWE corrected. BA – Broadmann area, k – cluster size, R – right hemisphere, L – left hemisphere, R – right hemisphere
